# Investigating the relationship between CRISPR-Cas content and growth rate in bacteria

**DOI:** 10.1101/2022.08.18.504381

**Authors:** Zhi-Ling Liu, En-Ze Hu, Deng-Ke Niu

**Affiliations:** MOE Key Laboratory for Biodiversity Science and Ecological Engineering and Beijing Key Laboratory of Gene Resource and Molecular Development, College of Life Sciences, Beijing Normal University, Beijing, 100875, China

**Keywords:** CRISPR-Cas, cell cycle, bacteria, prophage, trade-off

## Abstract

CRISPR-Cas systems provide adaptive immunity for prokaryotic cells by recognizing and eliminating the recurrent genetic invaders whose sequences had been captured in a prior infection and stored in the CRISPR arrays as spacers. However, the biological/environmental factors determining the efficiency of this immune system have yet to be fully characterized. Recent studies in cultured bacteria showed that slowing the growth rate of bacterial cells could promote their acquisition of novel spacers. This study examined the relationship between the CRISPR-Cas content and the minimal doubling time across the bacteria and the archaea domains. Every completely sequenced genome could be used to predict a minimal doubling time. With a large dataset of 4142 bacterial samples, we found that the predicted minimal doubling times are positively correlated with spacer number and other parameters of the CRISPR-Cas systems, like array number, *Cas* gene cluster number, and *Cas* gene number. Different datasets gave different results. Weak results were obtained in analyzing bacterial empirical minimal doubling times and the archaea domain. Still, the conclusion of more spacers in slowly-grown prokaryotes was supported. In addition, we found that the minimal doubling times are negatively correlated with the occurrence of prophages, and the spacer numbers per array are negatively associated with the number of prophages. These observations support the existence of an evolutionary trade-off between bacterial growth and adaptive defense against virulent phages.

## INTRODUCTION

CRISPR-Cas systems provide adaptive immunity by recognizing and eliminating the recurrent invaders whose genetic information had been captured in a prior infection and stored in the CRISPR arrays as spacers. Accumulating evidence indicates that slowing the host cell growth rate could enhance spacer acquisition, probably because the slow growth affects virus replication more than the cell’s response to the invaders. Using a fluorescent CRISPR adaptation reporter method, Amlinger et al. (1) quantified the adaptation efficiency of type II CRISPR in different growth phases of *Escherichia coli*. They found that spacer acquisition occurs predominantly during the late exponential/early stationary phase. In line with this study, McKenzie et al. (2) observed that the *E. coli* cells that adapted and cleared the target DNA earlier grew significantly slower than the population means. The soil-dwelling bacterium *Pseudomonas aeruginosa* can infect plant and human bodies. It grows slowly in the cooler temperatures of the soil or during plant infection and rapidly under the higher temperature of human bodies. Høyland-Kroghsbo et al. (3) found that decreasing the growth rate could increase the CRISPR adaptation frequency of the bacterium. Recently, Dimitriu et al. (4) found that the bacteriostatic agents that slow down the growth of *P. aeruginosa* could promote the phage-derived acquisition of novel spacers into the CRISPR array. Furthermore, they showed that limited carbon sources associated with slow bacterial growth could also enhance the evolution of CRISPR immunity. The latter result is in line with the previous observations of increased CRISPR-Cas immunity in bacteria grown in a nutrient-limited medium (5). Dormancy could be regarded as a prolonged growth state of the prokaryotic cells. In the response of the type VI CRISPR-Cas system to RNA virus infection, the Cas13 enzymes of the bacteria *Listeria ivanovii* were found to destruct both the RNA virus genomes and the host mRNA molecules, which further induce dormancy of the host bacterial cells (6). In the same study, the immunity of the bacterial cells against the unrelated virus was enhanced in the type-VI-CRISPR-Cas-system-induced dormant state.

All the above results were obtained from analyzing a few cultured bacteria in laboratories. We could deduce that environmental factors that drive the bacteria to grow slowly in nature would enhance CRISPR-Cas immunity. The present study examines whether there is a universal relationship between growth rate and adaptive immunity across the bacteria domain. The null hypothesis to be tested is that CRISPR-Cas contents are unrelated to the growth rates across the bacteria and archaea domains.

## RESULTS

### Construction of the datasets

We constructed three datasets of bacterial minimal doubling times and CRISPR-Cas system contents (Tables S1-S3). The first dataset contains 262 bacterial species with their empirical minimal doubling times retrieved from reference (7). Multiple strains have been sequenced for some bacterial species. In total, 424 genomes were downloaded from the GenBank database (8). These genomes have a wide variation in CRISPR-Cas contents (Table S4), on average, 1.34 CRISPR arrays (ranging from 0 to 12), 43.95 spacers (ranging from 0 to 665), 0.875 *Cas* gene clusters (ranging from 0 to 10), and 5.21 *Cas* genes (ranging from 0 to 49). Among them, 247 genomes do not have any CRISPR-Cas components, 20 have only CRISPR arrays or *Cas* gene clusters, and 157 have both CRISPR arrays and *Cas* gene clusters. For the species with multiple sequenced strains in this dataset, we averaged the multiple CRISPR-Cas content values of each species for further analysis.

The second dataset contains 4142 bacterial genomes with their predicted minimal doubling times obtained from the EGGO database (accessed Nov 26, 2021) (9).

In the third dataset, 508 bacterial genomes, each with only one CRISPR array and one *Cas* gene cluster, were selected from the second dataset.

To control the effects of growth temperature, we also integrated the optimal growth temperatures predicted by the program Tome (version 1.0.0) (10) in the three datasets. Each species’ predicted optimal growth temperatures in the first dataset were also averaged.

From the descriptive statistics of the three datasets (Table 1), we can see that the predicted minimal doubling times have a similar median value and a similar range but a lower mean value than the empirical minimal doubling times. Phylogenetic generalized least squares (PGLS) regression analysis showed that the empirical minimal doubling times and the predicted minimal doubling times are significantly correlated (*n* = 252, slope = 0.228, *P* = 1.5 × 10^-10^).

**Table 1.** Descriptive statistics for the minimal doubling times of all the bacteria in our datasets and the top 5% CRISPR-spacer-rich bacteria.

| Datasets | *n* | Median | Mean | Standard deviation | Maximum | Minimum |
| --- | --- | --- | --- | --- | --- | --- |
| The first dataset |  |  |  |  |  |  |
| All the bacteria | 262 | 2.66 | 6.71 | 8.70 | 52.80 | 0.23 |
| Top 5% | 13 | 6.50 | 11.57 | 10.62 | 36.94 | 0.92 |
| The second dataset |  |  |  |  |  |  |
| All the bacteria | 4142 | 2.30 | 3.41 | 3.24 | 54.17 | 0.12 |
| Top 5% | 207 | 6.571 | 6.66 | 4.24 | 18.84 | 0.19 |
| The third dataset |  |  |  |  |  |  |
| All the bacteria | 508 | 2.25 | 3.60 | 4.16 | 54.17 | 0.14 |
| Top 5% | 25 | 2.88 | 4.51 | 4.85 | 19.99 | 0.58 |

Please see the beginning of the Result section for the definition of the three datasets. We compared the minimal doubling times between the top 5% CRISPR-spacer-rich bacteria and the other 95% bacteria in the three datasets using phylANOVA (12). The F values (and *P* values) obtained in analyzing the three datasets are 4.3 (0.096), 231.3 (0.001), and 1.3 (0.356), respectively.

### Higher CRISPR-Cas contents in bacteria with long minimal doubling times

We calculated the descriptive statistics for the minimal doubling times of the bacteria with the top 5% highest spacer numbers (Table 1). From the descriptive statistics, it appears that the bacteria rich in spacers grow slower than other bacteria. However, phylogenetic ANOVA analysis did not find significant differences between the top 5% CRISPR-spacer-rich bacteria and the other 95% bacteria in the first or the third dataset (*P* > 0.05 for both cases, Table 1). We suspected that their small sample sizes might account for the insignificant results.

Previously, Vieira-Silva and Rocha (11) discretized the minimum doubling time *(D)* into four classes: very fast (*D* < 1h), fast (1h < *D* < 2h), intermediate (2h < *D* < 5h) and slow (*D* ≥ 5h). Referring to their classification, our dichotomy between fast and slow was repeated three times, with one h, two h, and five h as the boundary. We compared the slowly-grown bacteria and fast-grown bacteria using the phylogenetic ANOVA method, phyloANOVA (12), and found that the former had higher CRISPR-Cas contents, not just more spacers, than the latter in the second dataset when the slow was defined as *D* ≥ 5h and the fast was defined as *D* < 5h (Fig. 1A-D). Meanwhile, we noticed that the slowly-grown bacteria have significantly higher optimal growth temperatures than the fast-grown bacteria (Fig. 1E). Significant differences were not found in the other two datasets or the phylogenetic ANOVA analyses using other definitions of fast and slow in the second dataset. More results with significant differences were obtained when the CRISPR-Cas contents were compared using the Mann-Whitney *U* test. As ignoring the effects of common ancestry in evolutionary analysis often gave false-positive results (13, 14), we are inclined to accept the results of the phylogenetic ANOVA analysis.

**Fig. 1.**
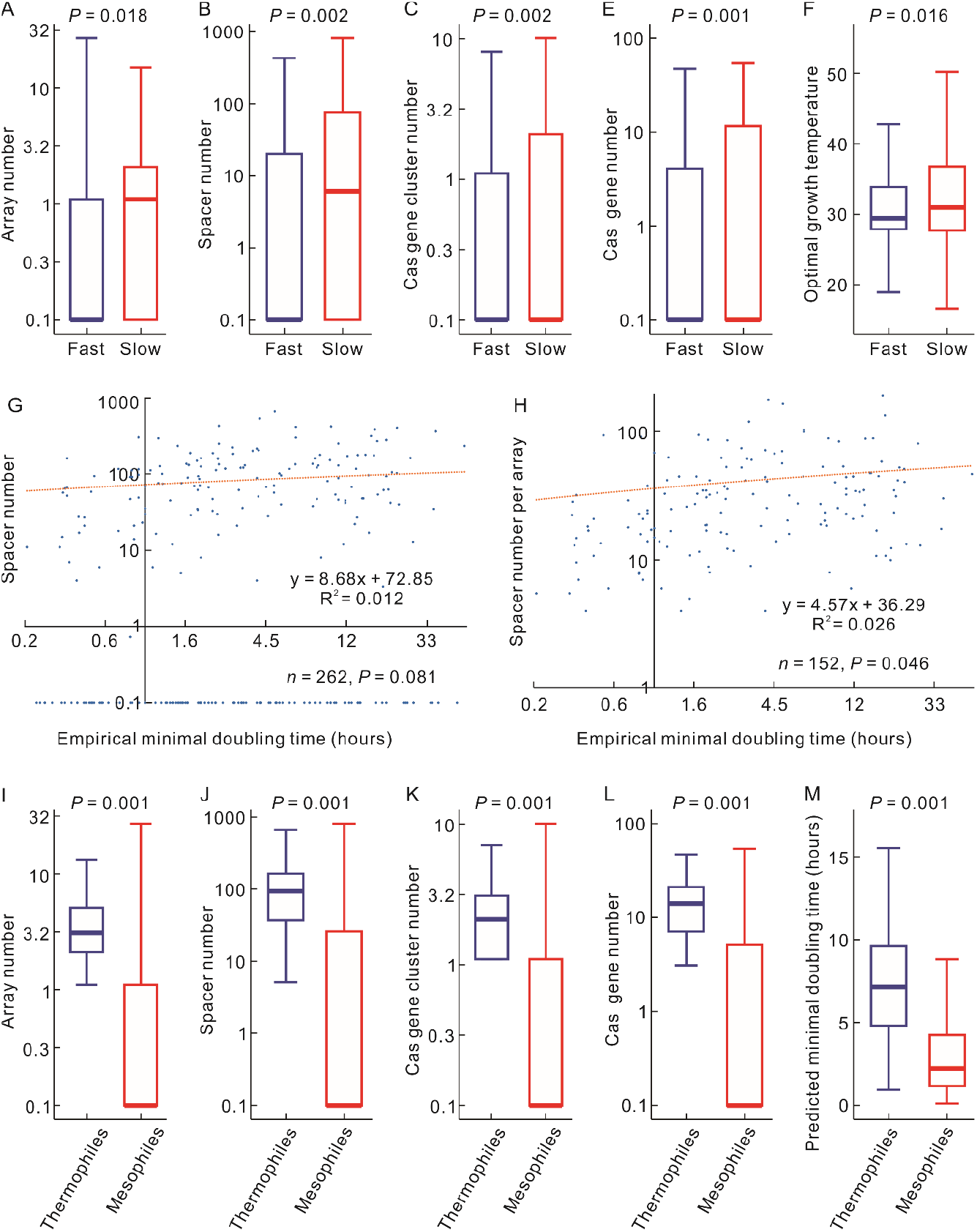
Correlations among bacterial CRISPR-Cas contents, the minimal doubling times, and optimal growth temperatures. (A-E) Slowly-grown bacteria (*n* = 891) have higher CRISPR-Cas contents and higher optimal growth temperatures than fast-grown bacteria (*n* = 3251). The slowly-grown was defined as the minimal doubling times ≥ 5h and the fast-grown was defined as the minimal doubling times < 5h. The comparisons were performed using phylANOVA with the *P* values adjusted by the Benjamini-Hochberg (BH) procedure, and the F values of A-E are 178.0, 293.5, 341.4, 365.7, and 186.6, respectively. (G) The CRISPR spacer numbers and (H) spacer numbers per array in bacteria are positively correlated with the empirical minimal doubling times. The correlations were examined using PGLS regression analyses, with the positive sign of the regression slope to indicate the positive correlation. Actual hours were labeled in these two panels just because they are intuitive. The doubling time values were log-transformed using base *e* in the PGLS analysis. (I-M) Thermophilic bacteria (*n* = 156) have higher CRISPR-Cas contents and longer doubling times than mesophilic bacteria (*n* = 3944). The comparisons were performed using phylANOVA with the *P* values adjusted by the Benjamini-Hochberg (BH) procedure, and the F values of A-E are 321.6, 474.9, 509.0, 634.0, and 241.8, respectively. Following reference (38), mesophiles were defined as 20 ≤ optimal growth temperature < 45 °C, and thermophiles were defined as 45 ≤ optimal growth temperature < 80 °C.

For a global relationship between CRISPR-Cas contents and growth rates across the bacterial domains, we performed PGLS regression analysis using CRISPR-Cas contents as the dependent variable and the minimal doubling times as the independent variable. The first dataset has a positive correlation at a marginally significant level (*n* = 262, slope = 8.683, *P* = 0.081. Fig. 1G and Table 2) between the empirical minimal doubling times and the spacer numbers. To control the effects of growth temperatures, we performed multiple PGLS regression analyses using optimal growth temperatures as the second independent variable. In this analysis, the empirical minimal doubling times are significantly correlated with spacer numbers (*n* = 262, slope = 13.145, *P* = 0.009), *Cas* gene cluster numbers (*n* = 262, slope = 0.166, *P* = 0.035), and *Cas* gene numbers (*n* = 262, slope = 0.985, *P* = 0.032) (Table 2).

**Table 2.** Correlations of the minimal doubling times *(D)* with CRISPR-Cas contents.

| | $N$ | $D$ | | $D$ with OPT controlled | | OPT with $D$ controlled | |
| --- | --- | --- | --- | --- | --- | --- | --- |
| | | Slope | $P$ | Slope | $P$ | Slope | $P$ |
| The first dataset |  |  |  |  |  |  |  |
| Array numbers | 262 | 0.035 | 0.791 | 0.138 | 0.285 | **0.056** | **4×10<sup>-5</sup>** |
| Spacer numbers | 262 | 8.683 | 0.081 | **13.145** | **0.009** | **2.187** | **4×10<sup>-4</sup>** |
| Spacer numbers per array | 152 | **4.570** | **0.046** | **5.047** | **0.033** | 0.183 | 0.439 |
| *Cas* gene cluster numbers | 262 | 0.096 | 0.235 | **0.166** | **0.035** | **0.039** | **3×10<sup>-6</sup>** |
| *Cas* gene numbers | 262 | 0.505 | 0.286 | **0.985** | **0.032** | **0.261** | **10<sup>-7</sup>** |
| *Cas* gene numbers per cluster | 149 | −0.003 | 0.977 | 0.066 | 0.552 | **0.025** | **0.010** |
| The second dataset |  |  |  |  |  |  |  |
| Array numbers | 4142 | **0.168** | **0.002** | **0.151** | **0.005** | **0.056** | **10<sup>-14</sup>** |
| Spacer numbers | 4142 | **5.326** | **0.003** | **5.212** | **0.003** | **1.456** | **2×10<sup>-8</sup>** |
| Spacer numbers per array | 1507 | **3.282** | **0.010** | **3.408** | **0.008** | −0.101 | 0.434 |
| *Cas* gene cluster numbers | 4142 | **0.091** | **0.001** | **0.085** | **0.002** | **0.033** | **10<sup>-17</sup>** |
| *Cas* gene numbers | 4142 | **0.562** | **9×10<sup>-4</sup>** | **0.528** | **0.001** | **0.203** | **3×10<sup>-18</sup>** |
| *Cas* gene numbers per cluster | 1077 | 0.106 | 0.145 | 0.105 | 0.153 | 6×10<sup>-4</sup> | 0.931 |
| The third dataset |  |  |  |  |  |  |  |
| Spacer numbers | 508 | **4.351** | **0.029** | **4.588** | **0.021** | -0.421 | 0.115 |
| *Cas* gene numbers | 508 | 0.177 | 0.107 | 0.172 | 0.118 | 0.010 | 0.515 |

OPT, optimal growth temperature. Please see the beginning of the Result section for the definition of the datasets. The correlations were examined using multiple PGLS regression analyses, with the positive sign of the regression slope to indicate the positive correlation. Significant, marginally significant, and nonsignificant results were shown in bold, regular, and grey, respectively

In the second dataset, all the CRISPR-Cas content parameters (array number, spacer number, *Cas* gene cluster number, and *Cas* gene number) have significant positive correlations with the predicted minimal doubling times, no matter whether the optimal growth temperatures were controlled or not (slope > 0, *P* < 0.05 for all cases, Table 2).

The increase of spacer number with the minimal doubling time can be explained by an enhancing effect on spacer acquisition of slow growth as previously observed in the cultured bacteria (1–6). However, slowly grown bacteria may also have more spacers because they have a more significant number of CRISPR arrays, either duplicated or acquired horizontally. The same logic exists for the *Cas* gene number. Therefore, we examined the relationship of minimal doubling times with the spacer number per array and *Cas* gene number per cluster (Here, only genomes with arrays or *Cas* gene clusters were counted. Genomes without CRISPR-Cas systems were not included in the denominator). The spacer numbers per array positively correlate with the empirical minimal doubling times (*n* = 152, slope = 4.570, *P* = 0.046. Fig. 1H) and the predicted minimal doubling times (*n* = 1507, slope = 3.282, *P* = 0.010). However, the *Cas* gene numbers per cluster are not significantly correlated with the empirical minimal doubling time (*n* = 149, slope = −0.003, *P* = 0.977) or the predicted minimal doubling time (*n* = 1472, slope = 0.106, *P* = 0.145). In addition, we also examined the relationship of predicted minimal doubling times with spacer numbers and *Cas* gene numbers in the third dataset, i.e., 508 bacterial genomes, each of which has only one CRISPR array and one *Cas* gene cluster. Only spacer numbers significantly correlate with the predicted minimal doubling times, regardless of whether optimal growth temperatures were controlled (Table 2).

According to the *Cas* genes (15), 508 bacteria in the third dataset could be classified, including 329 bacteria with class 1 *Cas* gene cluster (304 type I, 21 type III, and four type IV) and 173 bacteria with class 2 *Cas* gene cluster (167 type II, five type V, and one type VI). We performed the PGLS regression analyses for the groups with more than 50 samples, class 1, class 2, type I, and type II. Although the bacteria with type I *Cas* gene cluster is a subset of the bacteria with class 1 *Cas* gene cluster, they did not give consistent correlations. In bacteria with class 1 *Cas* gene cluster, the spacer numbers are marginally correlated with the minimal doubling times after controlling the optimal growth temperatures (*n* = 329, slope = 4.028, *P* = 0.097). However, the relationship is far from statistically significant in the bacteria with type I Cas gene cluster (*P* > 0.10). Significant (*P* < 0.05) or marginally significant (0.05 ≤ *P* < 0.1) correlations have not been observed in bacteria with class 2 or type II *Cas* gene clusters. Different types of CRISPR-Cas systems seem not identical in their relationships with minimal doubling times. Future studies with larger samples will be helpful for a conclusion.

### CRISPR-Cas content and growth temperature

Previous studies observed that bacterial CRISPR abundance is positively correlated with growth temperatures (16–19), with an abrupt jump at around 45°C (20). Here, we first compared the CRISPR-Cas contents and predicted minimal doubling times between thermophilic and mesophilic bacteria using the phylogenetic ANOVA method, phyloANOVA (12). As shown in Fig. 1I-M, thermophilic bacteria have higher CRISPR-Cas contents and longer doubling times. Therefore, we examined the relationship between the CRISPR-Cas contents and optimal growth temperatures by controlling the minimal doubling time using multiple PGLS regression analyses. The results show that the minimal doubling times and the growth temperatures are independently correlated with CRISPR-Cas contents (Table 2). However, the spacer numbers per array and the *Cas* gene numbers per cluster in the second dataset are not significantly correlated with the optimal growth temperatures *(P* > 0.10 for both cases, Table 2). Similarly, the spacer and *Cas* gene numbers in the third dataset are not significantly correlated with the optimal growth temperatures (*P* > 0.10 for both cases, Table 2). It seems that the increase of adaptive immunity with growth temperature is mainly achieved by increasing the number of CRISPR arrays and *Cas* gene clusters. By contrast, increasing adaptive immunity with minimal doubling time seems achieved by increasing all aspects of the CRISPR-Cas systems.

### Relationship between prophage contents, minimal doubling times, and CRISPR-Cas contents

Previously, Touchon et al. found that the frequency of prophage is negatively correlated with the minimal doubling time in bacteria (21). We examined this relationship using our datasets. Neither prophage numbers nor the presence/absence of prophages significantly correlate with bacterial empirical minimal doubling times (*P* > 0.10 for both cases). However, both prophage numbers and the presence/absence of prophages are negatively correlated with bacterial predicted minimal doubling times *(n* = 4142, slope = −0.186, *P* = 5 × 10^-5^ and *n* = 4142, slope = −0.057, *P* = 10^-5^, respectively). In the third dataset, the presence/absence of prophages is negatively correlated with the predicted minimal doubling time (*n* = 508, slope = −0.052, *P* = 0.040). Furthermore, we examined the relationship between prophage contents and CRISPR-Cas systems. The spacer numbers per array in the first and second datasets and the spacer numbers in the third dataset consistently correlate negatively with the prophage numbers (*P* < 0.05 for all the cases, Table 3). However, only the spacer numbers of the third dataset are negatively correlated (at a marginally significant level) with the prophage existence. Table 3 shows that prophages have weak or no relationship with the array numbers, the *Cas* gene cluster numbers, or the *Cas* gene numbers.

**Table 3.** Correlations between the prophage contents and CRISPR-Cas contents.

|  |  | Prophage number |  | Prophage existence |  |
| --- | --- | --- | --- | --- | --- |
| | $N$ | Slope | $P$ | Slope | $P$ |
| The first dataset |  |  |  |  |  |
| Array numbers | 262 | −0.025 | 0.606 | 0.003 | 0.783 |
| Spacer numbers | 262 | **−0.002** | **0.037** | $−3\times10^{-4}$ | 0.313 |
| Spacer numbers per array | 262 | **−0.010** | **0.007** | −0.001 | 0.268 |
| *Cas* gene cluster numbers | 262 | −0.062 | 0.422 | −0.010 | 0.633 |
| *Cas* gene numbers | 262 | −0.013 | 0.332 | −0.003 | 0.321 |
| *Cas* gene numbers per cluster | 262 | −0.085 | 0.316 | **−0.054** | **0.029** |
| The second dataset |  |  |  |  |  |
| Array numbers | 4142 | 0.017 | 0.256 | −0.002 | 0.621 |
| Spacer numbers | 4142 | $−3\times10^{-4}$ | 0.421 | $−10^{-4}$ | 0.216 |
| Spacer numbers per array | 4142 | **−0.005** | **0.002** | −0.001 | 0.160 |
| *Cas* gene cluster numbers | 4142 | 0.039 | 0.169 | −0.001 | 0.856 |
| *Cas* gene numbers | 4142 | 0.005 | 0.341 | −0.001 | 0.608 |
| *Cas* gene numbers per cluster | 4142 | −0.028 | 0.284 | −0.005 | 0.448 |
| The third dataset |  |  |  |  |  |
| Spacer numbers | 508 | **−0.005** | **0.009** | **−0.001** | **0.093** |
| *Cas* gene numbers | 508 | −0.016 | 0.672 | 0.002 | 0.837 |

The correlations were examined using PGLS regression analyses, with the negative sign of the regression slope to indicate the negative correlation. The statistically significant (and marginally significant) cases were shown in bold. Prophage existence: 0 and 1 were designated as absence and presence in the PGLS analysis. Please see the beginning of the
Result section for the definition of the three datasets.

## DISCUSSION

In a few cultured bacterial species, slow growth has been found to enhance spacer acquisition (1–6). Our phylogenetic analyses showed that the spacer numbers and the other CRISPR-Cas content parameters (array number, *Cas* gene cluster number, and *Cas* gene number) positively correlate with the minimal doubling times in bacteria.

Previous authors reporting the enhancing effects of slow growth on spacer acquisition generally explain their observation by the mechanisms of spacer acquisition (1, 3, 4). If the slow growth could enhance spacer acquisition, the CRISPR-Cas systems would be an efficient mechanism to counteract viruses in slowly-grown cells, and so be selected when the growth rate has been slowed down on an evolutionary scale.

Furthermore, we could put the CRISPR-Cas-enhancing effect of slow growth into an ecological context. Studies on natural bacterial communities support the existence of trade-offs between intraspecific competition ability (faster growth) and invulnerability to predation (including virus resistance) (22–25). With the limited resources available in natural environments, the bacteria that allocate more resources to defense have to grow slowly, and the bacteria with high competition ability allocate less amount of resources to virus resistance. The trade-off between faster growth and defense is destined if maintaining and expressing the CRISPR-Cas system exert a significant metabolic burden. This metabolic burden has been observed in *Streptococcus thermophilus* by comparing a strain that constitutively expresses the Cas protein and a strain with a non-functional mutant of Cas (26). However, acquiring a few spacers was not associated with fitness costs (26). Besides the metabolic burden, autoimmunity induced by self- and prophage-targeting spacers is another widely-cited cost of CRISPR-Cas systems (27, 28). A trade-off between fast growth and defense using CRISPR-Cas systems might be mediated by the autoimmunity targeting prophages. In the present study, we found evidence for the mutually exclusive effects between prophage and CRISPR-Cas immunity across the bacterial domain (Table 3). Because prophages are more frequent in bacteria with short minimal doubling times (21), there should be a selective force against CRISPR–Cas systems in fast-grown bacteria (29, 30).

In this study, the three bacterial datasets did not always give consistent results. We suggest that the differences in the results should be attributed to the differences in the accuracy and the datasets’ sample size. In a previous study, we emphasized that phylogenetic comparative studies generally required larger samples than typical statistical analyses (31). Here, we want to briefly discuss the empirical and predicted data’s advantages and disadvantages. At first glance, the empirical minimal doubling times seem superior to the predicted minimal doubling times because the prediction methods could not achieve an accuracy of 100%. As the wide-spread diversity among the strains designated as one bacterial species (32), the ideal empirical data should include different parameters (e.g., minimal doubling time, optimal growth temperature, and CRISPR-Cas contents) collected from the same strain. Unfortunately, this is not always the case. In many cases, a phenotype value is assigned to a bacterial species without mentioning the strain identity, so the values of different phenotypes might come from different strains. By contrast, all the predicted phenotype values of a strain come from the genome sequences of the same strain. Potential errors resulting from polymorphism within each species could be eliminated if all the parameters were predicted/annotated from genome sequences.

We also examined the relationships in archaea using PGLS regression. First, we found that the predicted minimal doubling times are not significantly correlated with the empirical minimal doubling times (*n* = 100, slope = 0.055, *P* = 0.325). The method to predict minimal doubling times was mainly based on bacterial data (9). It seems not to work accurately in archaea. Therefore, we only examined the relationship between archaea’s empirical minimal doubling times and CRISPR-Cas contents. Univariate PGLS regression did not find significant correlations (*P* > 0.10 for all the cases). However, after controlling optimal growth temperatures, the empirical minimal doubling times are significantly correlated with the array numbers (*n* = 103, slope = 0.838, *P* = 0.027), spacer numbers (*n* = 103, slope = 23.239, *P* = 0.022), *Cas* gene cluster numbers (*n* = 103, slope = 0.378, *P* = 0.005), and *Cas* gene numbers (*n* = 103, slope = 2.120, *P* = 0.017). Meanwhile, the spacer numbers per array positively correlate with the empirical minimal doubling time (*n* = 88, slope = 7.476, *P* = 0.018). Because of the limited number of samples, we did not analyze the archaea in such detail as the bacteria.

## MATERIALS AND METHODS

The datasets of prokaryotic empirical minimal doubling times and predicted minimal doubling times were retrieved from reference (7) and the EGGO database (accessed Nov 26, 2021) (9), respectively. To avoid the outlier effects on the statistical results, we only retained the minimal doubling time shorter than 60h. Because the data on minimal doubling times are highly skewed, they were log-transformed using base *e*. To cover more prokaryotes, we used the optimal growth temperatures predicted by the program Tome (version 1.0.0) (10). The phylogenetic trees and taxonomic information of bacteria and archaea were retrieved from the Genome Taxonomy Database (GTDB, accessed Apr 8, 2022) (33). Because each prokaryotic species rather than each strain has one empirical minimal doubling time value, phylogenetic comparative analysis of the empirical minimal doubling times requires a species tree. From the GTDB, we constructed a species tree using the representative genome of each species. The prokaryotic genome assembly files were downloaded from the GenBank database (accessed Apr 26, 2021) (8). The CRISPR-Cas systems of the prokaryotic genomes were annotated using CRISPRCasFinder (version 4.2.20) (34). Using VirSorter 2.2.3 (35), prophages were found in bacteria but not archaeal genomes. To avoid underestimation of the CRISPR-Cas systems, partially assembled genomes have been discarded, and only those with assembly levels of “complete genome” or “chromosome” were retained in our analyses. The *Cas* gene number of each type of *Cas* gene cluster is relatively constant in evolution, but different types and classes have a significant difference in their *Cas* gene number, ranging from 1 to 11 (15). The difference in *Cas* gene number between genomes should be attributed to different types of *Cas* gene clusters.

We estimated the phylogenetic signals (λ) of the analyzed characters using the R (Version 4.0.2) package phytools (Version 0.7-47) (36). All the analyzed characters exhibit significant phylogenetic signals, indicating that phylogenetic comparative methods are required in the analyses to control the effect of common ancestry. The correlations between different characters were estimated by the PGLS regression using the R (Version 4.0.2) package phylolm (version 2.6.2) (37). Pagel’s λ model has been applied in this study. The phylogenetic ANOVA analyses were performed using phyloANOVA (12) with the *P* values adjusted by the Benjamini-Hochberg (BH) procedure.

## Supporting information

All the supplementary tables.

## SUPPLEMENTAL MATERIAL

Supplemental material is available online only.

SUPPLEMENTAL FILE 1, XLSX file, 0.674 MB.

## ACKNOWLEDGMENTS

This work was supported by the National Natural Science Foundation of China (grant number 31671321). We thank the anonymous reviewers for their constructive comments.

## Notes

### Competing Interest Statement

The authors have declared no competing interest.

### Summary of Updates

More results have been presented in the figure.

## REFERENCES

1. Amlinger L, Hoekzema M, Wagner EGH, Koskiniemi S, Lundgren M. 2017. Fluorescent CRISPR Adaptation Reporter for rapid quantification of spacer acquisition. Scientific Reports 7:10392.

2. McKenzie RE, Keizer EM, Vink JNA, van Lopik J, Büke F, Kalkman V, Fleck C, Tans SJ, Brouns SJJ. 2022. Single cell variability of CRISPR-Cas interference and adaptation. Molecular Systems Biology 18:e10680.

3. Høyland-Kroghsbo NM, Muñoz KA, Bassler BL. 2018. Temperature, by controlling growth rate, regulates CRISPRCas activity in *Pseudomonas aeruginosa*. mBio 9:e02184–18.

4. Dimitriu T, Kurilovich E, Lapinska U, Severinov K, Pagliara S, Szczelkun MD, Westra ER. 2022. Bacteriostatic antibiotics promote CRISPR-Cas adaptive immunity by enabling increased spacer acquisition. Cell host & microbe 30:1–10.

5. Westra ER, van Houte S, Oyesiku-Blakemore S, Makin B, Broniewski JM, Best A, Bondy-Denomy J, Davidson A, Boots M, Buckling A. 2015. Parasite exposure drives selective evolution of constitutive versus inducible defense. Current Biology 25:1043–1049.

6. Meeske AJ, Nakandakari-Higa S, Marraffini LA. 2019. Cas13-induced cellular dormancy prevents the rise of CRISPR-resistant bacteriophage. Nature 570:241–245.

7. Madin JS, Nielsen DA, Brbic M, Corkrey R, Danko D, Edwards K, Engqvist MKM, Fierer N, Geoghegan JL, Gillings M, Kyrpides NC, Litchman E, Mason CE, Moore L, Nielsen SL, Paulsen IT, Price ND, Reddy TBK, Richards MA, Rocha EPC, Schmidt TM, Shaaban H, Shukla M, Supek F, Tetu SG, Vieira-Silva S, Wattam AR, Westfall DA, Westoby M. 2020. A synthesis of bacterial and archaeal phenotypic trait data. Scientific Data 7:170.

8. Sayers EW, Cavanaugh M, Clark K, Pruitt KD, Schoch CL, Sherry Stephen T, Karsch-Mizrachi I. 2021. GenBank. Nucleic Acids Research 50:D161–D164.

9. Weissman JL, Hou S, Fuhrman JA. 2021. Estimating maximal microbial growth rates from cultures, metagenomes, and single cells via codon usage patterns. Proceedings of the National Academy of Sciences 118:e2016810118.

10. Li G, Rabe KS, Nielsen J, Engqvist MKM. 2019. Machine learning applied to predicting microorganism growth temperatures and enzyme catalytic optima. ACS Synthetic Biology 8:1411–1420.

11. Vieira-Silva S, Rocha EPC. 2010. The systemic imprint of growth and its uses in ecological (meta)genomics. PLOS Genetics 6:e1000808.

12. Garland T, Jr., Dickerman AW, Janis CM, Jones JA. 1993. Phylogenetic analysis of covariance by computer simulation. Systematic Biology 42:265–292.

13. Felsenstein J. 1985. Phylogenies and the comparative method. The American Naturalist 125:1–15.

14. Symonds MRE, Blomberg SP. 2014. A primer on phylogenetic generalised least squares, p 105–130. In Garamszegi LZ (ed), Modern Phylogenetic Comparative Methods and Their Application in Evolutionary Biology: Concepts and Practice doi:10.1007/978-3-662-43550-2_5. Springer Berlin Heidelberg, Berlin, Heidelberg.

15. Makarova KS, Wolf YI, Iranzo J, Shmakov SA, Alkhnbashi OS, Brouns SJJ, Charpentier E, Cheng D, Haft DH, Horvath P, Moineau S, Mojica FJM, Scott D, Shah SA, Siksnys V, Terns MP, Venclovas C, White MF, Yakunin AF, Yan W, Zhang F, Garrett RA, Backofen R, van der Oost J, Barrangou R, Koonin EV. 2020. Evolutionary classification of CRISPR-Cas systems: a burst of class 2 and derived variants. Nature Reviews Microbiology 18:67–83.

16. Makarova KS, Wolf YI, Snir S, Koonin EV. 2011. Defense islands in bacterial and archaeal genomes and prediction of novel defense systems. Journal of Bacteriology 193:6039–6056.

17. Anderson RE, Brazelton WJ, Baross JA. 2011. Using CRISPRs as a metagenomic tool to identify microbial hosts of a diffuse flow hydrothermal vent viral assemblage. FEMS Microbiology Ecology 77:120–133.

18. Weinberger AD, Wolf YI, Lobkovsky AE, Gilmore MS, Koonin EV. 2012. Viral diversity threshold for adaptive immunity in prokaryotes. mBio 3:e00456–12.

19. Gophna U, Kristensen DM, Wolf YI, Popa O, Drevet C, Koonin EV. 2015. No evidence of inhibition of horizontal gene transfer by CRISPR–Cas on evolutionary timescales. The ISME Journal 9:2021–2027.

20. Lan X-R, Liu Z-L, Niu D-K. 2022. Precipitous increase of bacterial CRISPR-Cas abundance at around 45°C. Frontiers in Microbiology 13:773114.

21. Touchon M, Bernheim A, Rocha EPC. 2016. Genetic and life-history traits associated with the distribution of prophages in bacteria. The ISME Journal 10:2744–2754.

22. Cram JA, Parada AE, Fuhrman JA. 2016. Dilution reveals how viral lysis and grazing shape microbial communities. Limnology and Oceanography 61:889–905.

23. Yang JW, Chang F-H, Yeh Y-C, Tsai A-Y, Chiang K-P, Shiah F-K, Gong G-C, Hsieh C-h. 2021. Trade-off between competition ability and invulnerability to predation in marine microbes; protist grazing versus viral lysis effects. BioRxiv doi:10.1101/2021.10.09.463787.

24. Winter C, Bouvier T, Weinbauer MG, Thingstad TF. 2010. Trade-offs between competition and defense specialists among unicellular planktonic organisms: the “killing the winner” hypothesis revisited. Microbiology and Molecular Biology Reviews 74:42–57.

25. Våge S, Bratbak G, Egge J, Heldal M, Larsen A, Norland S, Lund Paulsen M, Pree B, Sandaa R-A, Skjoldal EF, Tsagaraki TM, Øvreås L, Thingstad TF. 2018. Simple models combining competition, defence and resource availability have broad implications in pelagic microbial food webs. Ecology Letters 21:1440–1452.

26. Vale PF, Lafforgue G, Gatchitch F, Gardan R, Moineau S, Gandon S. 2015. Costs of CRISPR-Cas-mediated resistance in *Streptococcus thermophilus*. Proceedings of the Royal Society B: Biological Sciences 282:20151270.

27. Bernheim A. 2017. [Why so rare if so essential: the determinants of the sparse distribution of CRISPR-Cas systems in bacterial genomes]. Biologie Aujourdhui 211:255–264.

28. Zaayman M, Wheatley RM. 2022. Fitness costs of CRISPR-Cas systems in bacteria. Microbiology 168:001209.

29. Rollie C, Chevallereau A, Watson BNJ, Chyou T-y, Fradet O, McLeod I, Fineran PC, Brown CM, Gandon S, Westra ER. 2020. Targeting of temperate phages drives loss of type I CRISPR–Cas systems. Nature 578:149–153.

30. Nobrega FL, Walinga H, Dutilh BE, Brouns SJJ. 2020. Prophages are associated with extensive CRISPR–Cas auto-immunity. Nucleic Acids Research 48:12074–12084.

31. Hu E-Z, Lan X-R, Liu Z-L, Gao J, Niu D-K. 2022. A positive correlation between GC content and growth temperature in prokaryotes. BMC Genomics 23:110.

32. Cobo-Simón M, Hart R, Ochman H. 2023. *Escherichia coli:* What is and which are? Molecular Biology and Evolution 40:msac273.

33. Parks DH, Chuvochina M, Rinke C, Mussig AJ, Chaumeil P-A, Hugenholtz P. 2021. GTDB: an ongoing census of bacterial and archaeal diversity through a phylogenetically consistent, rank normalized and complete genome-based taxonomy. Nucleic Acids Research 50:D785–D794.

34. Couvin D, Bernheim A, Toffano-Nioche C, Touchon M, Michalik J, Néron B, Rocha EPC, Vergnaud G, Gautheret D, Pourcel C. 2018. CRISPRCasFinder, an update of CRISRFinder, includes a portable version, enhanced performance and integrates search for Cas proteins. Nucleic Acids Research 46:W246–W251.

35. Guo J, Bolduc B, Zayed AA, Varsani A, Dominguez-Huerta G, Delmont TO, Pratama AA, Gazitúa MC, Vik D, Sullivan MB, Roux S. 2021. VirSorter2: a multi-classifier, expert-guided approach to detect diverse DNA and RNA viruses. Microbiome 9:37.

36. Revell LJ. 2012. phytools: an R package for phylogenetic comparative biology (and other things). Methods in Ecology and Evolution 3:217–223.

37. Ho LST, Ane C. 2014. A linear-time algorithm for Gaussian and non-Gaussian trait evolution models. Systematic Biology 63:397–408.

38. Sato Y, Okano K, Kimura H, Honda K. 2020. TEMPURA: database of growth TEMPeratures of Usual and RAre Prokaryotes. Microbes and Environments 35:ME20074.

